# 2 MV/cm Pulsed Electric Fields Promote Transthyretin Amyloid Disintegration

**DOI:** 10.1101/2020.02.15.950501

**Authors:** Gen Urabe, Takashi Sato, Gomaru Nakamura, Yoshihiro Kobashigawa, Hiroshi Morioka, Sunao Katsuki

## Abstract

Exposing transthyretin amyloid to 1000 sub-nanosecond 2 MV/cm pulsed electric fields (PEFs) promotes both amyloid disassembly and amyloid-derived transthyretin disintegration. The process produced no change in pH, and the resulting temperature increase was less than 1 °C. We conclude that PEFs’ physical effects facilitate amyloid disassembly, rather than thermal or chemical effects, and provoke amyloid-derived transthyretin disintegration, the latter of which is reported here for the first time.

## Introduction

Various biological reactions to electric and pulsed electric fields (PEFs) have been reported.^1–6^ Although much discussion has focused on membrane dynamics and damage, attention is shifting to proteins because, both theoretically and experimentally, membranes and their associated proteins, such as channels, pumps, actin cables, and microtubules, respond to electric fields.^7–12^ However, most previous studies analyzed proteins in cells^13–15^ and, therefore, did not consider cellular activities’ contributions under electric fields. For example, protein phosphorylation in PEFs may be the result of trans-membrane calcium influx, which activates calcineurin and promotes kinase activation.^16^ To examine direct electrical effects, purified proteins must be exposed to PEFs, but membrane and membrane-associated proteins may be theoretically under the field of 1 MV/cm or more, derived from the equation (1). If an applied electric field (*E*) is 10 kV/cm, cell radius (*r*) is 10 μm, *θ* = 0, and membrane thickness is 10 nm, the electric field in the membrane (*E_m_*) is

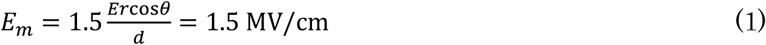

Because it is technically difficult to apply strong electric fields to liquid samples, most previous studies have used numerical calculations.^17–19^ According to such calculations, proteins change their three-dimensional structures under a static electric field of 1 MV/cm. Therefore, we focused on amyloid destruction theory, which speculates that amyloid is destroyed under fields stronger than 1 MV/cm which last longer than 400 ns.^20^ Many biologists and biochemists are skeptical about this theory because it is difficult to prove experimentally, and amyloid collapse has yet to be confirmed. However, verifying this theory would constitute a significant advance in biology, biochemistry, and bioelectrics. Pandey et al. presented results to support the theory but applied a weaker electric field of 230 kV/cm for longer than 40 hours. They used electrodes covered by Teflon and polydimethylsiloxane to prevent breakdown, both of which weaken electric fields in solution.^21^ In this paper, we describe a sub-nanosecond high-voltage generator and a specially designed chamber with a parallel gold-coated 1 mm electrode gap. We investigated PEFs’ thermal, chemical, and physical effects by applying 1000 pulses at 2 MV/cm to transthyretin amyloid.

## Results

### Transthyretin amyloid exposed to PEF

Transthyretin formed amyloid proteins at a pH of 4 and 37 °C (Fig. 1A). Our generator produced a voltage waveform of 200 kV with a 1 ns duration. A current emerged during voltage application, suggesting the sample reacted as a resistive and capacitive load. The electrode distance was 1 mm, making the maximum electric field 2 MV/cm (Fig. 1B). Application of 1000 pulses of a PEF at 2 MV/cm appeared to erase transthyretin amyloid of both the wild-type (WT) and L55P mutants, supporting the amyloid destruction theory (Fig. 1C, D, and E).

**Figure 1.**
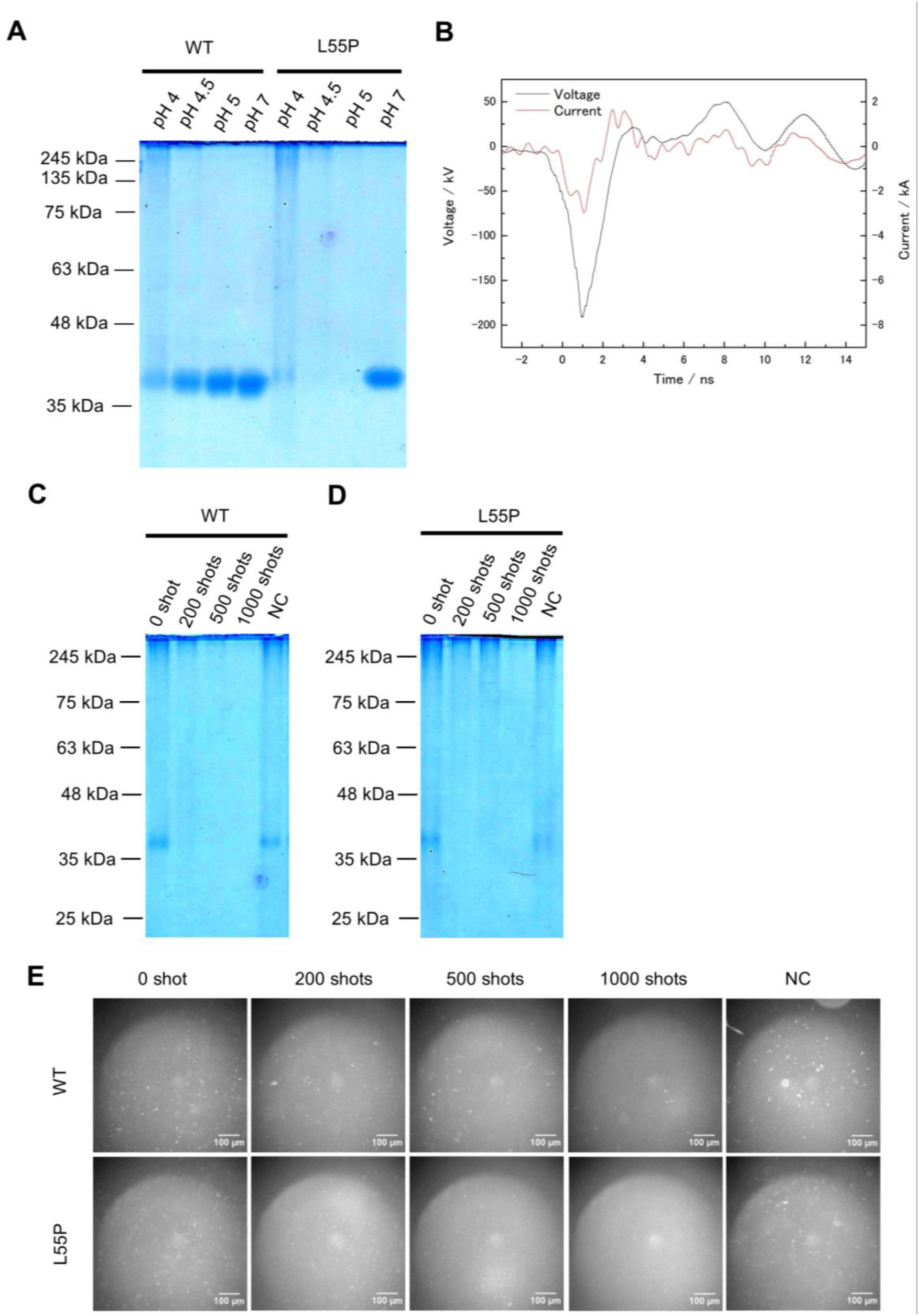
Transthyretin amyloid disappeared after being exposed to 1000 pulses at 2 MV/cm. **A**: Transthyretin of WT and L55P mutants formed amyloids with molar weights greater than 245 kDa at pH4 and 37 °C for 3 days. Transthyretin concentrations were 0.2 mg/mL. **B**: Voltage and current waveform. Both of them are the average of 20 waveformes. The maximum voltage on this graph was 200 kV and the maximum current was 3.5 kA. Amyloid of WT (**C**) and L55P mutant (**D**) disappeared after being exposed to 1000 pulses of a PEF at 2 MV/cm. Amyloid concentrations during PEF treatment were 0.2 mg/mL. NC was an amyloid solution (0.4 mg/mL) mixed with PEF-treated triple-diluted HEPES buffer at a volume ratio of 1:1. A, C, D were native PAGE and 1 μg of amyloid was in the gels. **E**: Amyloid fluorescent photos with a GFP filter. Application of 1000 pulses at 2 MV/cm reduced fluorescent dots. The white scale bar represents 100 μm. Each experiment was performed more than twice.

### Effects of reactive oxygen species, liquid pH, and liquid temperature can be ignored

Since PEF generates reactive oxygen species (ROS), mainly H_2_O_2_ in liquid samples,^22^ we measured the amount of H_2_O_2_ that emerged after 2 pulses of 2 MV/cm PEFs, detecting 0.08 ±0.01 μM H_2_O_2_ (Table 1). Assuming that the amount of H_2_O_2_ varies by pulse number, each concentration would be less than the pulse number times 0.08 ±0.01 μM. Amyloid did not collapse at any H_2_O_2_ concentration, even though at twice more than Figure 2A and 2B (Fig. 2A, B, C, and D). A comparison of the liquid pH before and after the application of 1000 pulses revealed no change from the initial value of 6.8 (Table 1). Furthermore, 1000 pulses raised the liquid temperature by less than 1 °C (Fig, 2E). This indicates direct electrical stress, rather than chemical or thermal effects, induced amyloid destruction.

**Figure 2.**
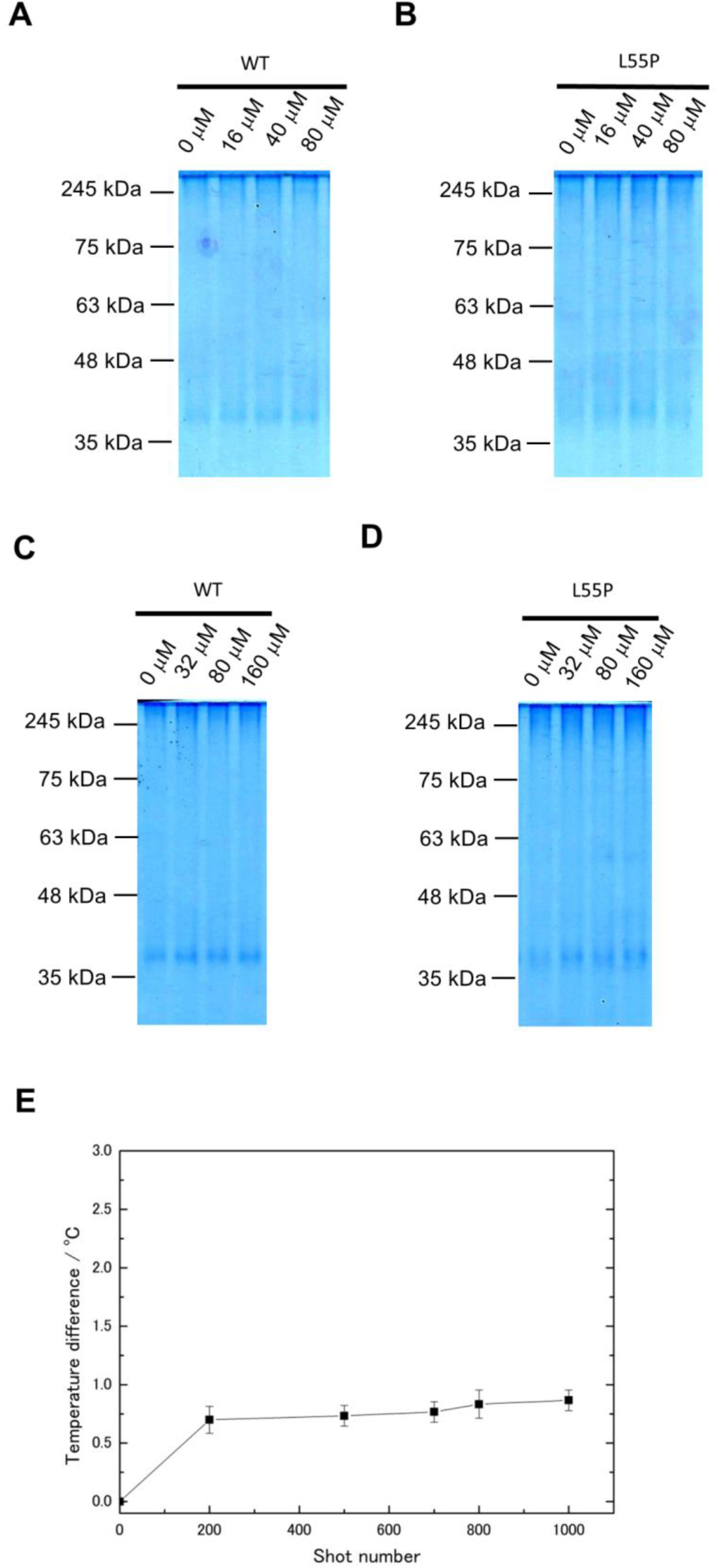
Consideration of typical chemical effects. H_2_O_2_ did not affect transthyretin amyloids on either WT (**A**, **C**) or L55P (**B**, **D**). Two pulses at 2 MV/cm generated 0.08 ±0.01 μM of H_2_O_2_. Concentrations in A and B were pulse number times 0.08 ±0.01 μM. That of C and D were twice the values of A and B. Each experiment was done once, but L55P at 160 μM was done twice. A, B, C, and D were native PAGE and 1 μg of amyloid was in the gels. All amyloid concentrations during H_2_O_2_ treatment were 0.2 mg/mL. **E**: Temperature rise at each pulse timing. The temperature increase at 1000 pulses was less than 1 °C. N = 3 and error bars were SEM.

**Table 1.**
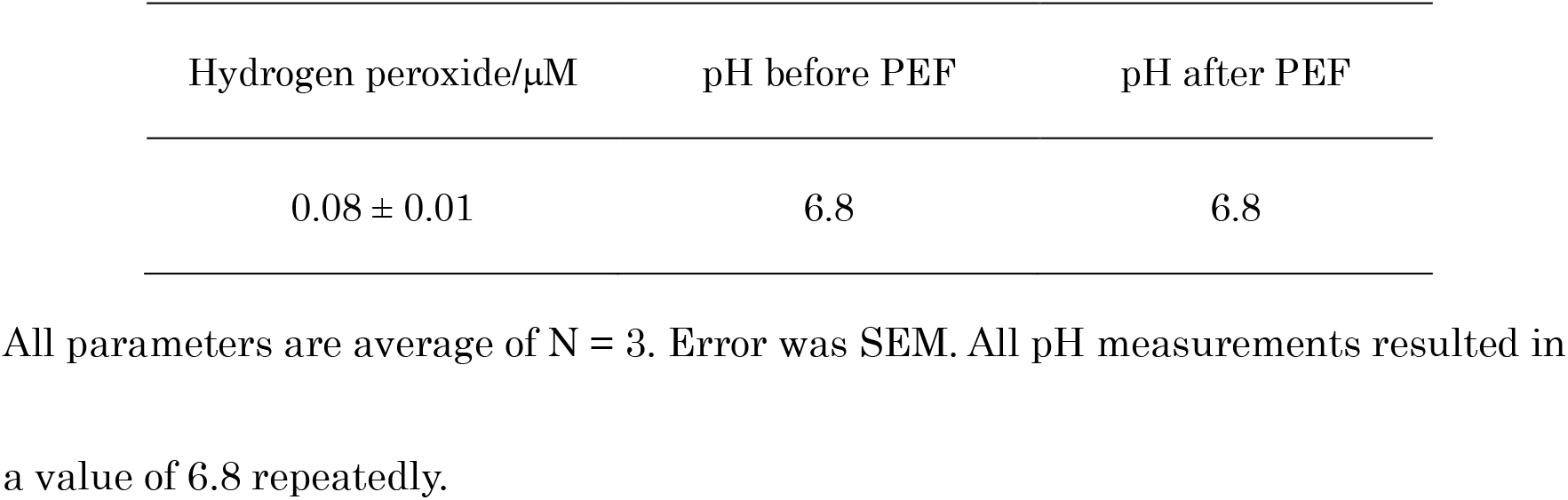
The amount of hydrogen peroxide that emerged after 2 pulses at 2 MV/cm and solution pH before and after PEF treatment.

### Stronger electric fields promoted amyloid disassembly

Exposing amyloid to 1000 pulses of a 2 MV/cm PEF destroyed amyloid more effectively than 1 MV/cm pulses, in a manner consistent with the proposed theory (Fig. 3A, B, and C).

**Figure 3.**
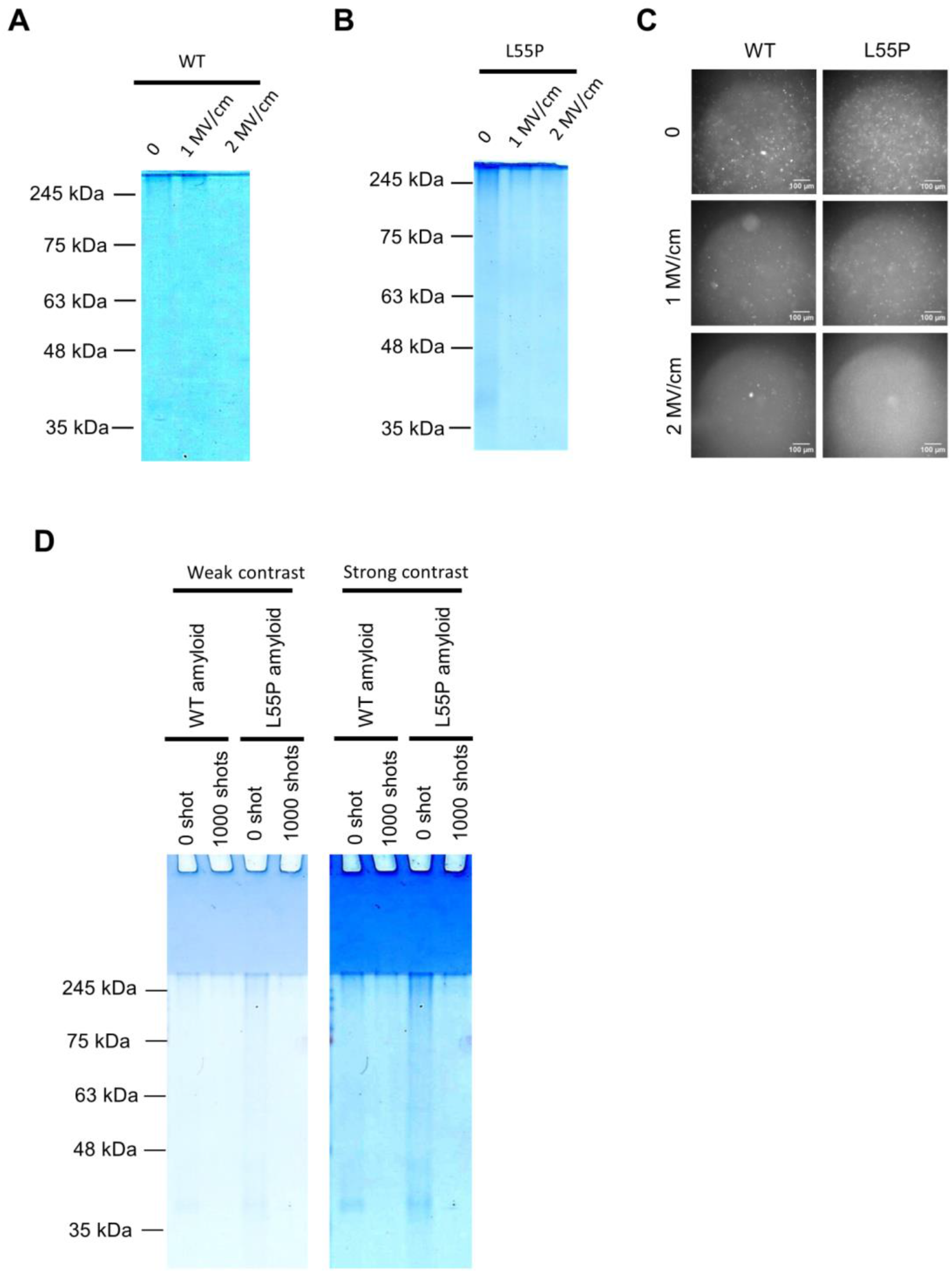
Effects of electric fields strength and the possibility of PEF-promoted aggregation. A weaker electric field (1 MV/cm) was less effective than 2 MV/cm PEF in both WT (**A**) and L55P (**B**). This tendency was as the same as in fluorescent analysis with a GFP filter (**C**). The pulses number was 1000 in A, B, and C. **D**: If proteins aggregated into a large molecule, strong bands emerged on the well bottom of the stacking gel. Such a signal did not emerge, although amyloid disappeared. A, B,C, were done twice but D once. A, B, and D were native PAGE and 1 μg of amyloid were in the gels. Concentrations of amyloids during PEF treatment were 0.2 mg/mL.

An electric field has the potential to form new covalent bonds in egg whites.^23–25^ A 2 MV/cm PEF may promote aggregation of amyloid with a molar weight too large to detect by gel electrophoresis, consequently reducing the amyloid band. A stacking gel photograph showed heavy molecules did not aggregate after 1000 pulses of 2 MV/cm PEF, supporting the amyloid disassemble hypothesis (Fig. 3D).

### PEF digested amyloid-derived transthyretin but not tetramer-derived transthyretin

To examine whether transthyretin persisted after PEF application, we performed reducing-condition sodium dodecyl sulfate-polyacrylamide gel electrophoresis (SDS-PAGE). The subunit band of transthyretin disappeared after 1000 pulses of a 2 MV/cm PEF in both WT and L55P mutants, indicating that amyloid transthyretin was digested (Fig. 4A and B). However, normal tetramer transthyretin was less affected than amyloid transthyretin; the tetramer transthyretin monomer band did not dramatically disappear at 1000 pulses of a 2 MV/cm PEF in either WT or L55P (Fig. 4C). Although non-specific bands between 63 kDa and 75 kDa usually appear in reduced SDS-PAGE, signal intensity and subunit band disappearance exhibited no apparent correlation (Supplementary Fig.4).

**Figure 4.**
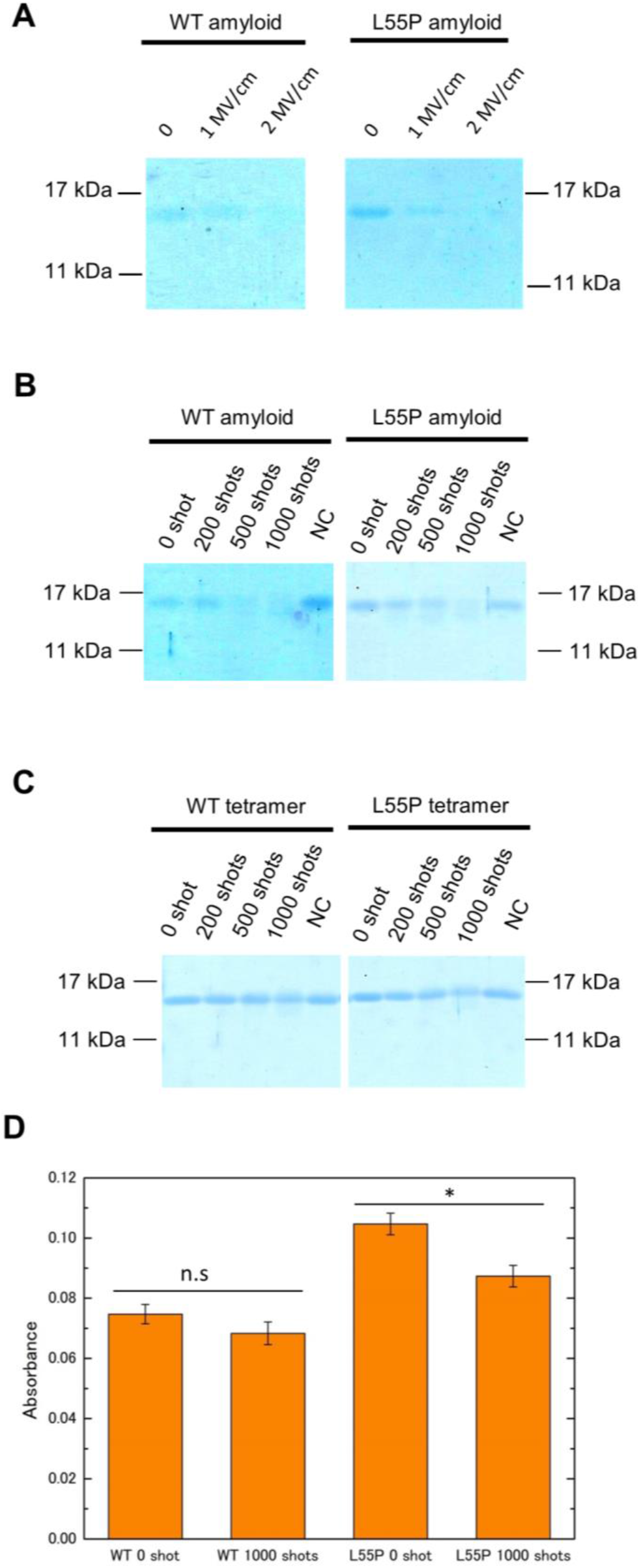
Transthyretin subunit analysis with SDS-PAGE. Application of 1000 pulses at 2 MV/cm destroyed amyloid-derived transthyretin subunit and weaker electric fields (**A**), while fewer pulses decreased the ability (**B**) in both WT and L55P mutants. The amyloid concentration during PEF treatment was 0.2 mg/mL in A and B. **C**: However, 1000 pulses at 2 MV/cm slightly reduced the tetramer-derived subunit band. The tetramer concentration during PEF treatment was 0.1 mg/mL in C. A, B, and C was performed twice with SDS-PAGE. A 0.35 μg sample of transthyretin was in the gels. Comparing the absorbance of amyloids using BCA, the L55P mutant exhibited a significant decrease after 1000 pulses at 2 MV/cm but WT proteins did not (n.s: no significant, *: p < 0.01, N = 3) (**D**).

Measurement of absorbance using the bicinchoninic acid (BCA), which detects peptide bonds, revealed that the absorbance of L55P amyloid with 1000 pulses at 2 MV/cm decreased significantly, suggesting that the PEF broke peptide bonds to digest transthyretin. The WT protein was not associated with significant reductions, but the treated samples tended to reduce signal strength (Fig. 4D).

## Discussion

### Amyloid destruction theory requires improvement

Our current results support the amyloid destruction theory but also suggest the theory should be modified to explain protein digestion, which had not been predicted and which did not happen in tetramer transthyretin. The differences between amyloid and tetramer transthyretin in SDS-PAGE can be explained by charge density derived from charged and polarized amino-acids, and mechanical oscillation frequency. The amyloid structure appeared much denser than the tetramers’ structure, implying that amyloid may be subject to greater electrical stress due to higher charge density.^26,27^ In terms of oscillation frequency, supramolecular vibration (whole-structure vibration) frequencies in microtubules and filaments cover the MHz-to-GHz range. However, molecular vibrations of normal proteins derived from weak bonds and atoms around the bonds are in the THz level.^28,29^ The PEFs’ main frequencies were near 50 MHz and 250 MHz, which was consistent with supramolecular vibrations (Fig. 5A, and 5B). PEFs of 2 MV/cm lasting 1 ns may efficiently promote sympathetic amyloid vibrations and, consequently, induce collapse of covalent bonds of transthyretin. To examine this hypothesis, we anticipate measuring impedance frequency characteristics of tetramer and amyloid transthyretin, adjusting the PEF duration to longer than 2 ns, in future experiments.

**Figure 5.**
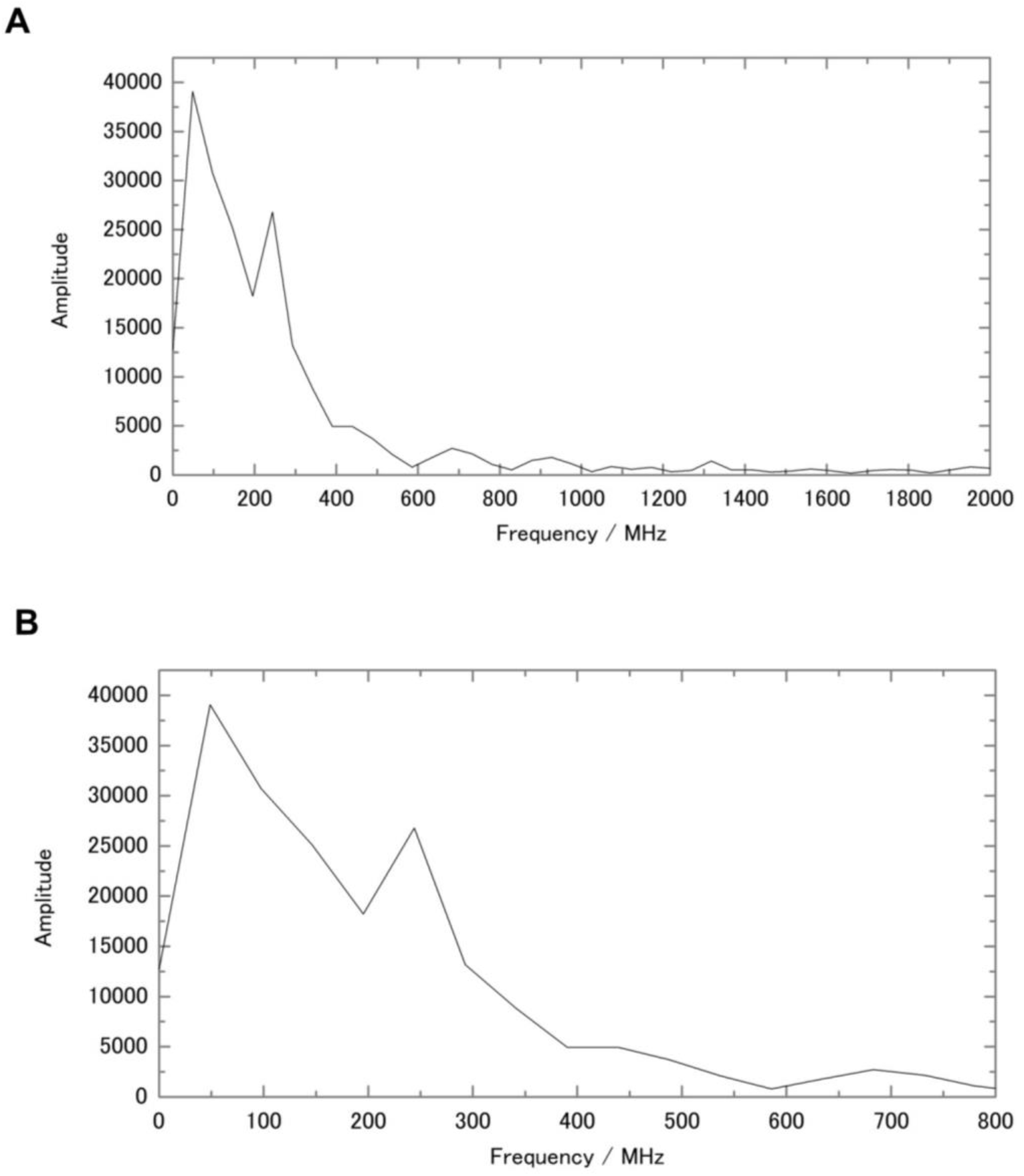
Fourier transformation of voltage in Figure 1B. **A**: Frequency distribution from 0 to 2000 MHz. **B**: Zooming up the range from 0 to 800 MHz, the PEF has two peaks at 50 MHz and 250 MHz.

One of the possible reasons WT amyloid was not significantly reduced (Figure 4D) is that the amount of WT amyloid was less than that of L55P (Fig. 1A and 3D), and the amount of decrease was too small to fulfill a criterion of significant decrease.

### Local pH shift on electrode surface vicinity

Non-buffer solutions can cause pH values to jump on a cathode surface and plunge on the anode side, while a buffer solution produces smaller effects.^30–32^ Transthyretin amyloid formed at pH 4 (Fig. 1A), amyloids at pH 9 remained (Fig. 1E and 3C), and amyloid was found in a HEPES buffer. These results suggest that a pH shift may not affect amyloid disintegration, regardless of whether a local pH shift occurs.

### Electric field emerges in sample solution

Some electrochemists consider voltage division of the applied PEF, due to electric bilayers on the cathode and anode, as parallel to resistance and capacitance. However, in high-frequency electric fields more than 1 MHz, the impedance of electric bilayers theoretically drops to zero, and the impedance of load will be almost identical to that of the solution.^33–36^ A 1 ns pulse width of 2 MV/cm has peaks at 50 MHz and 250 MHz frequency, and we believe that the current PEF was fully applied to the protein solutions.

### Conclusion

This paper not only supports the amyloid disassemble theory that electric fields stronger than 1 MV/cm promote amyloid collapse, but also suggests a need to add a protein digestion hypothesis to the theory. Our experiment confirmed that amyloid disintegration was caused by the physical effects of PEF, not the chemical or thermal effects.

## Materials

### Transthyretin preparation and amyloid formation

WT and L55P mutant transthyretin were expressed in *Escherichia coli* and then purified. A 0.4 mg/mL sample of transthyretin in HEPES buffer (50 mM HEPES, 150 mM NaCl, pH 6.8) was mixed with an acetate buffer (200 mM sodium acetate, 50 mM NaCl, pH 4) at a ratio of 20:20 μL and kept at 37 °C for 3 days. The pH 4.5 and pH 5 acetate buffers had the same composition, but a pH 7 solution was composed of a 50 mM NaCl solution. A 40 μL sample of amyloid was centrifuged at 20,000 *g* at room temperature for 5 minutes. After removing the supernatants, sediment amyloids were dissolved in a triple-diluted HEPES buffer (17 mM HEPES, 50 mM NaCl, pH 6.8) for pulse application.

### 2 MV/cm PEF generation

We used a nanosecond pulse to prevent plasma formation in the PEF exposure chamber. Our high-voltage nanosecond pulse generator consisted of a spark gap–driven 10-stage Marx circuit with an output capacitance of 94 pF; a pulse-peaking section, including a coaxial capacitor with the capacitance of 12 pF and a spark gap as the output switch; and another spark gap as a tail-cut switch for reducing the pulse to 1 ns. The Marx circuit was pressurized with 0.57 MPa nitrogen (N_2_) gas, whereas the pulse-peaking section and the tail-cut switch were pressurized with 0.5 MPa sulfur hexafluoride (SF_6_) gas to energize the system. We used a capacitive divider and a self-integrated pick-up coil to measure the voltage and current in the PEF exposure chamber, respectively. High-voltage pulses up to 200 kV and lasting 1 ns were delivered to 1 mm-gap, parallel-plane electrodes to generate an electric field exceeding 1 MV/cm. The electrodes were made of stainless steel (SUS316), and the anodic electrode was gold-coated to minimize the chemical effects of metal ions from the electrodes. The pulse repetition rate was fixed at 1.7 Hz to prevent the solution temperature from rising during PEF exposure.

### H_2_O_2_ detection, pH measurement, and temperature measurement

We used an Amplite Fluorimetric Hydrogen Peroxide Assay Kit *Near Infrared Fluorescence* from CosmoBio to measure H_2_O_2_ concentrations. We prepared a working solution (Amplite IR Peroxide Substrate, 0.8 U/mL peroxidase) and mixed the solution with miliQ at a ratio of 100:100 μL. The mixed solution was exposed to 2 MV/cm for 2 pulses and collected. Because the maximum treatable amount was 25 μL, we combined 9 samples of 2 pulse-solutions to prepare 200 μL of treated sample. A Quantus fluorometer measured the sample fluorescence with a red fluorescence filter to calculate the concentration of H_2_O_2_.

A LAQUAtwin compact pH meter was used to measure the pH of pre-treated and post-treated liquids. Because at least 100 μL is required for the measurements, we prepared 5 samples of 1000 pulse-solutions for the 100 μL treated samples.

An AMOTH FL-2400 fiberoptic thermometer with a FS300-2M probe measured on-time temperature of pulse-treated solution.

### Native PAGE

Native PAGE used a 4% stacking gel (4 w/v% acrylamide/bis mixed solution 29:1, 0.125 M Tris-Cl pH 6.8, 0.09 w/v%ammonium peroxodisulfate solution, 0.08 v/v% N,N,N′,N′-tetramethylethylenediamine) and 14 % separation gel (15 w/v% acrylamide/bis mixed solution 29:1, 0.375 M Tris-Cl pH 8.8, 0.091 w/v% ammonium peroxodisulfate solution, 0.08 v/v% N,N,N′,N′-tetramethylethylenediamine). Then, 5.5 μL of transthyretin solution and 5.5 μL of the sample buffer (0.1 M Tris-Cl pH 6.8, 20 v/v% glycerol, 0.05 w/v% bromophenol blue BPB) were mixed, and 10 μL of mixed solutions were applied to the wells of the gel. A Mini300 electric power source obtained from AS ONE applied a constant 15 mA current for 200 min for electrophoresis.

### Sodium dodecyl sulfate-polyacrylamide gel electrophoresis

SDS-PAGE used 4% stacking gel (4 w/v% acrylamide/bis mixed solution 29:1, 0.125 M Tris-Cl pH 6.8, 0.1 w/v% SDS, 0.09 w/v% ammonium peroxodisulfate solution, 0.08 v/v% N,N,N′,N ′-tetramethylethylenediamine) and 15% separation gel (15 w/v% acrylamide/bis mixed solution 29:1, 0.375 M Tris-Cl pH 8.8, 0.1 w/v% SDS, 0.091 w/v% ammonium peroxodisulfate solution, 0.08 v/v% N,N,N′,N′-tetramethylethylenediamine). Transthyretin solution and a sample buffer (0.1 M Tris-Cl pH 6.8, 20 v/v% glycerol, 4 w/v% SDS, 12 v/v% 2-mercaptoethanol, 0.05 w/v% BPB) were mixed at ratio of 1:1 and heated at 100 °C for 10 minutes. After cooling down the samples, they were centrifuged at 20,000 *g* for 1 min. Next, 3.5 μL of mixed solutions were applied to wells of the gel. A Mini300 electric power source obtained from AS ONE applied a constant 18 mA current for 110 min for electrophoresis.

### Amyloid fluorescent analysis

We used a fluorescent microscope (Leica, DMi8), combined with a digital camera (Canon, EOS 8000D), to observe amyloid fluorescence. Amyloid (0.1 mg/mL) was dissolved in thioflavin-T-glycine buffer (25 mM glycine, 10 μM thioflavin-T) and incubated on ice for 3 minutes without light. A green fluorescence protein filter was used.

### BCA method

This study used a protein assay BCA kit from Wako to measure the amount of transthyretin amyloid. An iMark microplate reader from BIO-RAD measured absorbance at 570 nm. Amyloid absorbance, without PEF application, was calculated by subtracting the raw absorbance of 0-pulse amyloid from that of a no-pulse triple-diluted HEPES buffer. Absorbance of 1000 pulses at 2 MV/cm was acquired by subtracting the raw absorbance of treated amyloid from that of exposed triple-diluted HEPES buffer.

### Statistical analysis

The data were presented as the mean ± standard error (SEM) for n samples (as shown in Fig. 5). Statistical analyses were performed using a two-tailed t-test; p < 0.01 was considered statistically significant.

## Acknowledgments

This study was partially supported by a Grant-in-Aid for Scientific Research (17H03220). The authors would like to thank Enago (www.enago.jp) for the English-language review.

## Authors’ contribution

S.K. and G.U. built the pulse generator, measured the voltage and current, and designed the research. G.N. provided assistance building the pulse generator. T.S, Y.K and H.M. offered advice on protocol and purified transthyretin to operate this project.

---

**Supplementary Figure 1.**
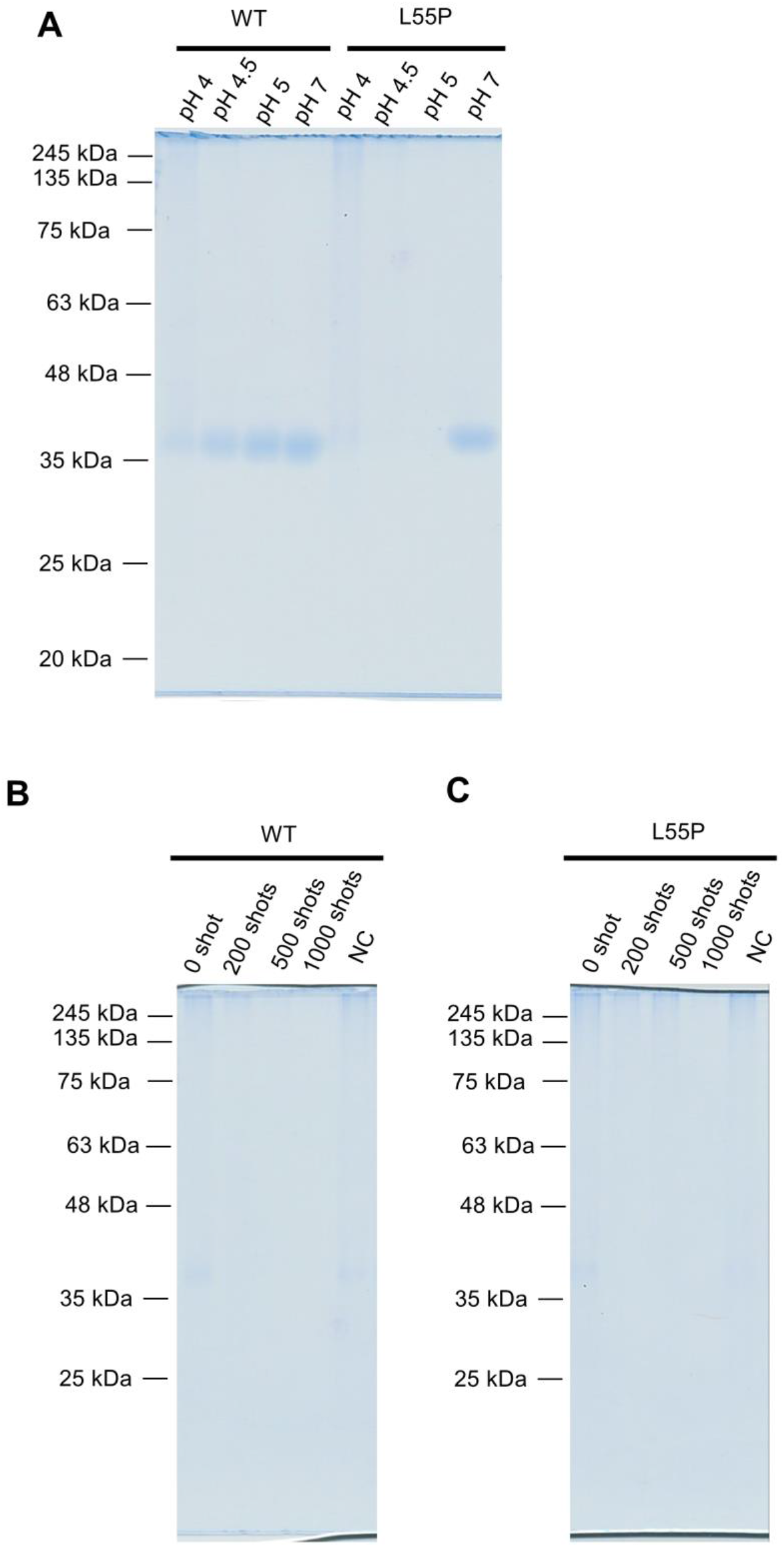
Full photographs without contrast of gel electrophoresis in Figure 1. **A**: Full photo without contrast of amyloid formation of Figure 1A. **B** and **C**: Full photographs without contrast of Figure 1C and 1D.

**Supplementary Figure 2.**
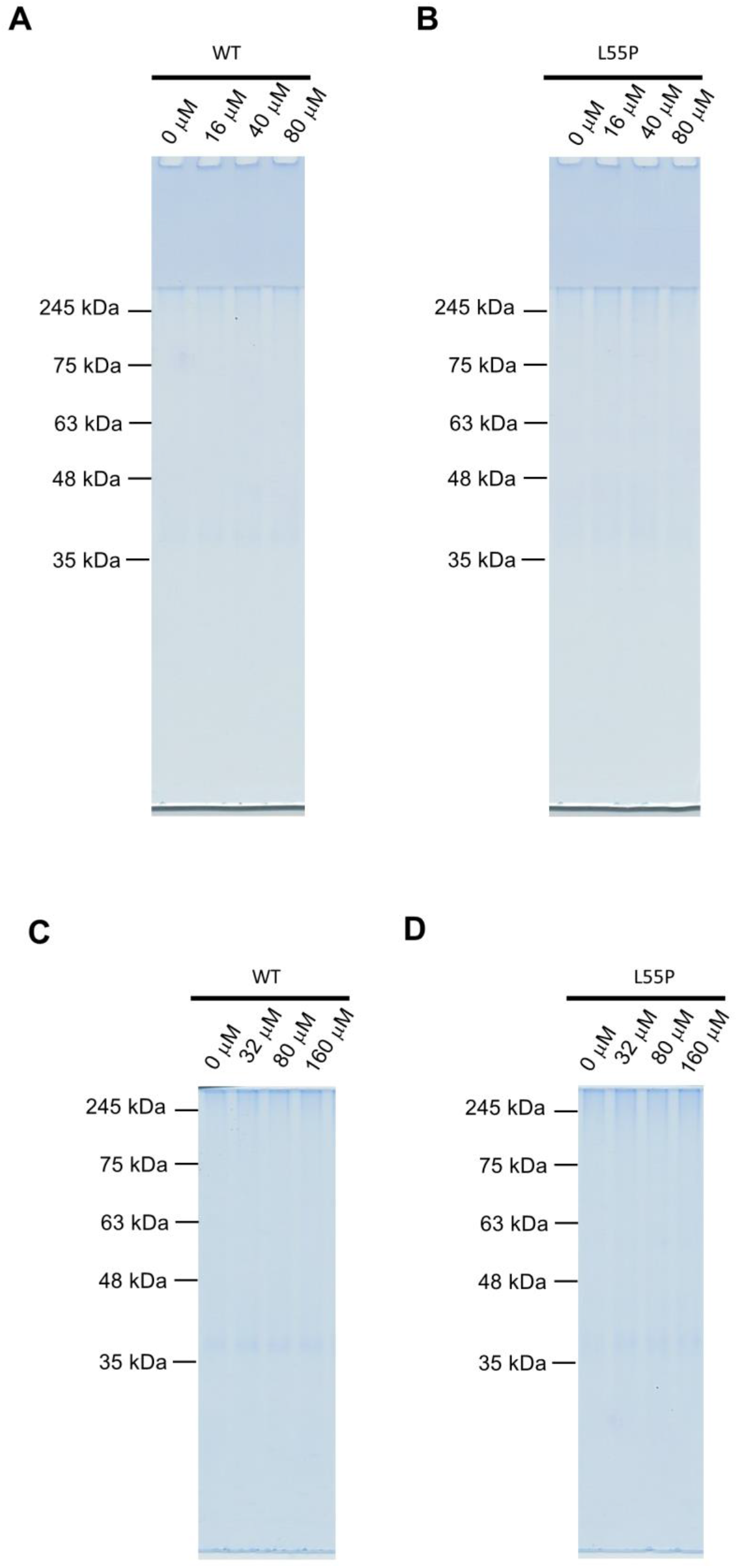
Full photographs without contrast of gel electrophoresis in Figure 2. All characters for tags are the same as those in Figure 2.

**Supplementary Figure 3.**
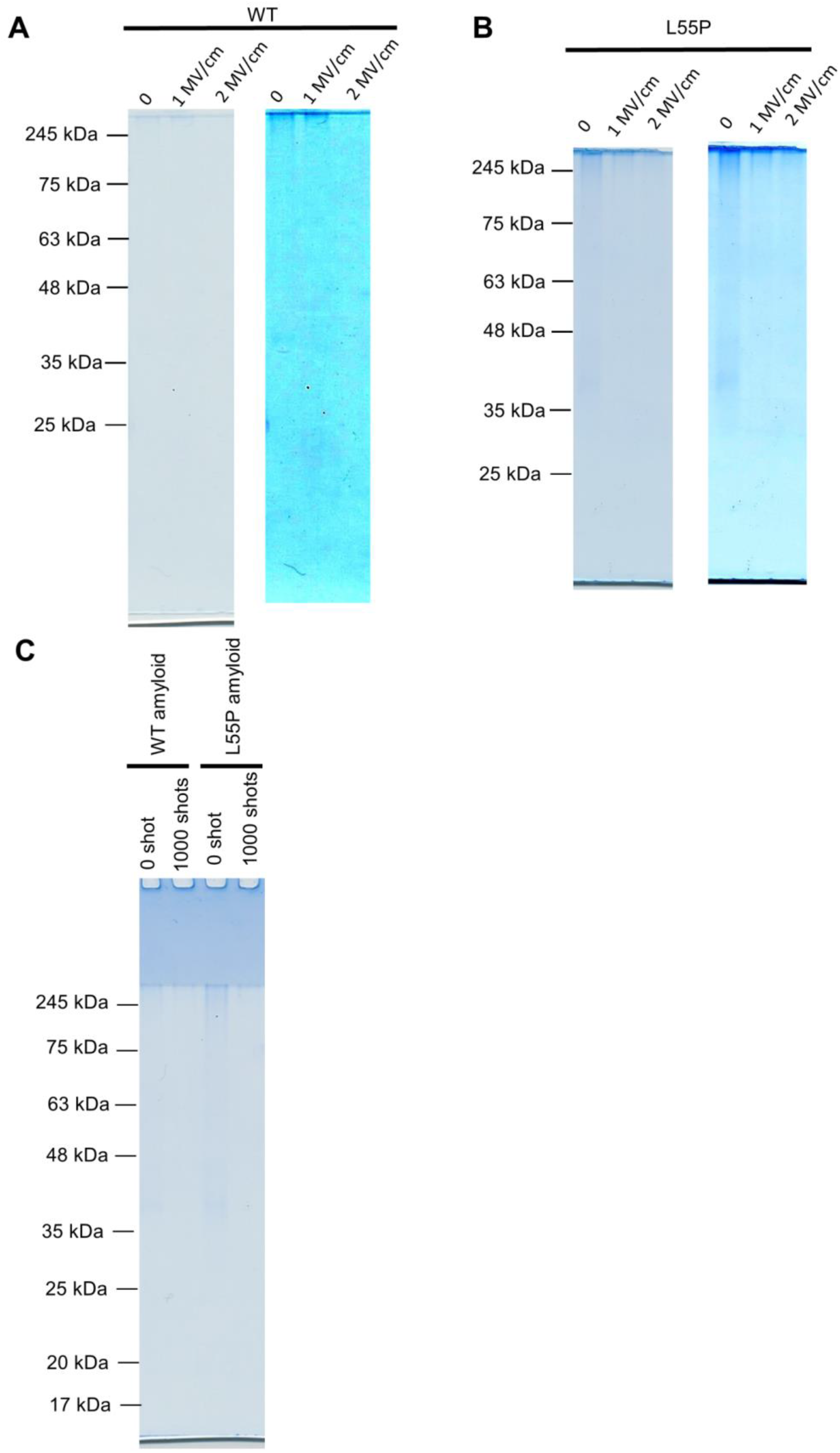
Full photographs of gel electrophoresis in Figure 3. **A** and **B**: Full photos of Figure 3A and 3B. Because the bands were too weak to check without contrast, contrast versions were produced. **C**: Full photograph without contrast of Figure 3D.

**Supplementary Figure 4.**
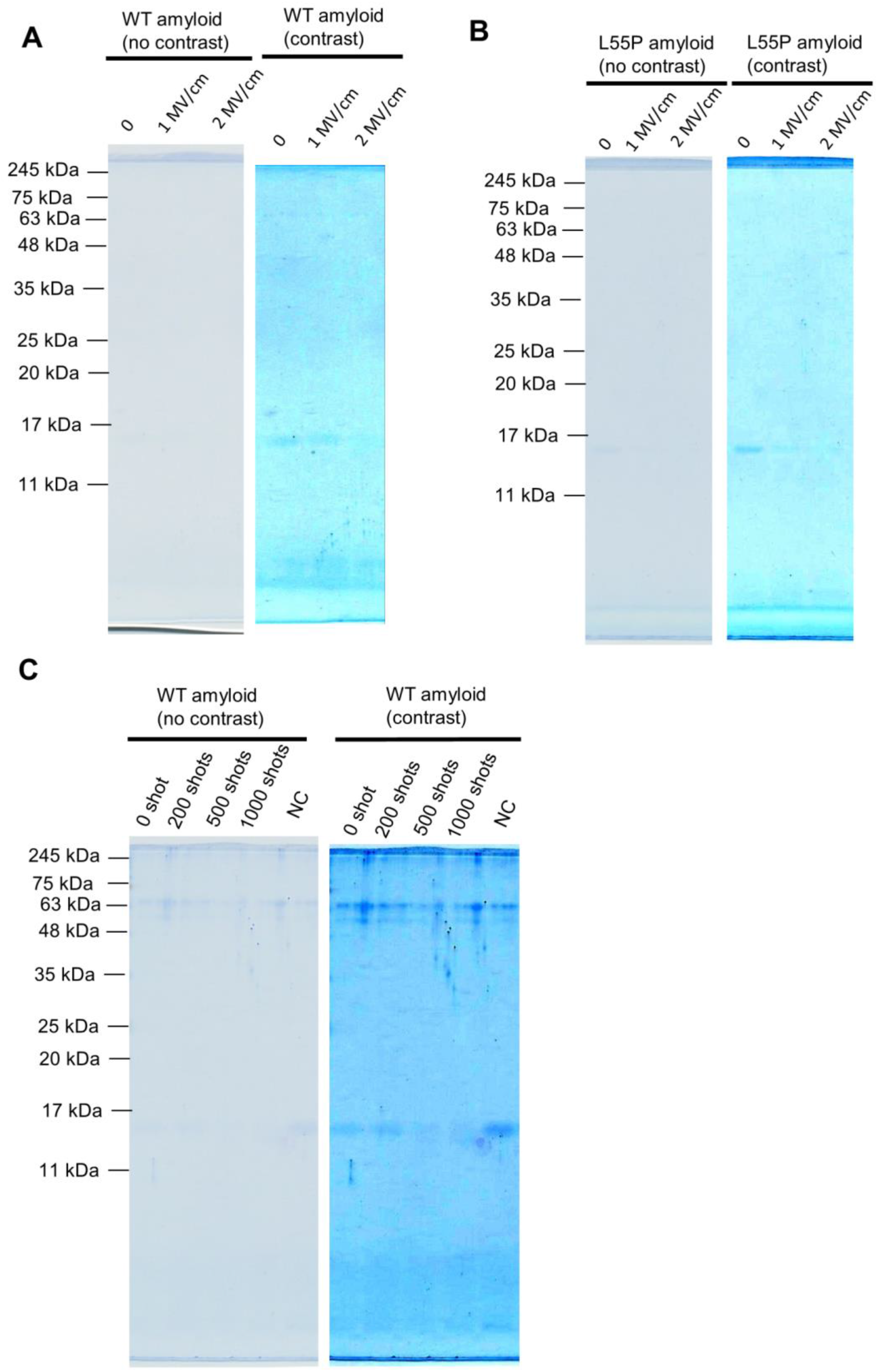
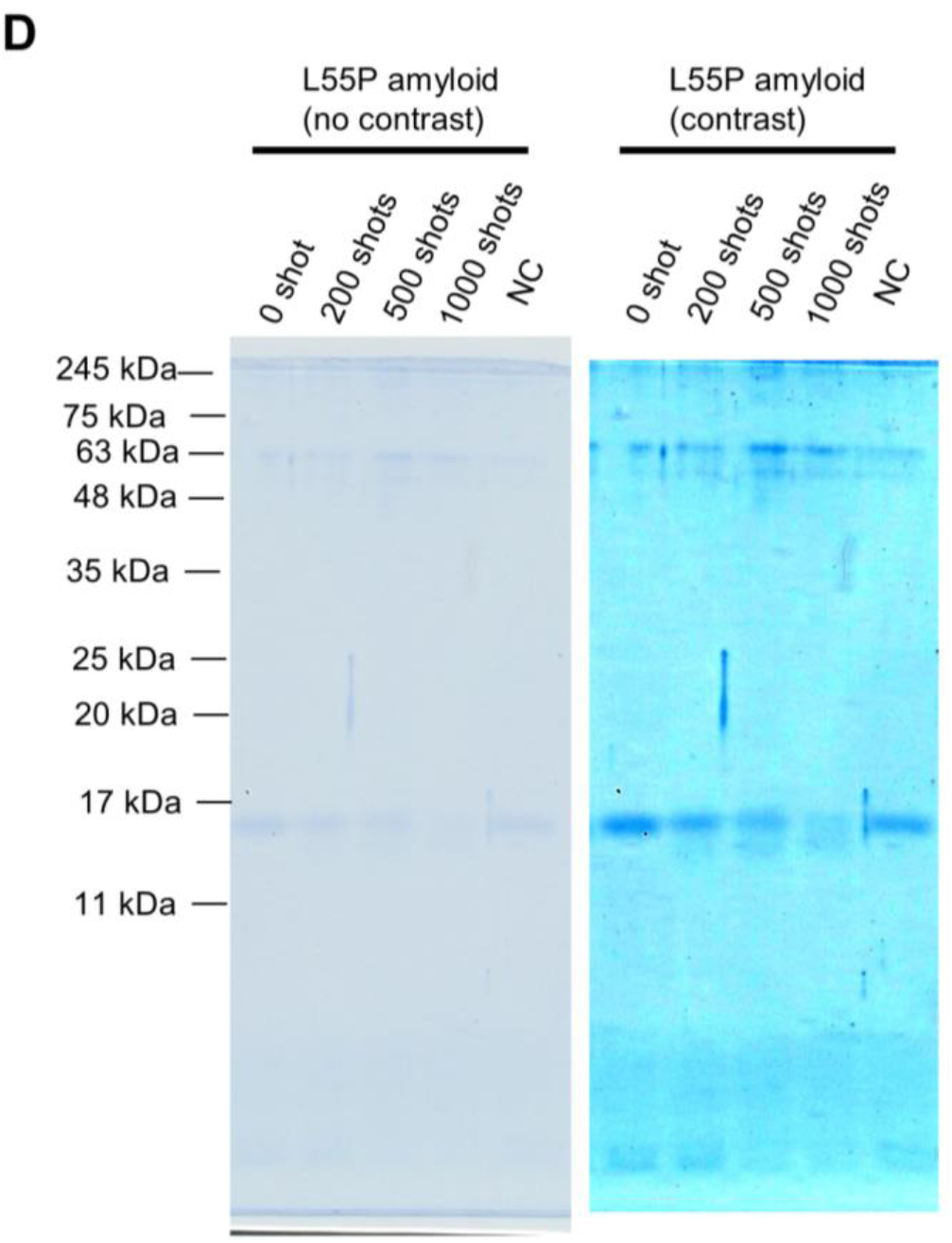
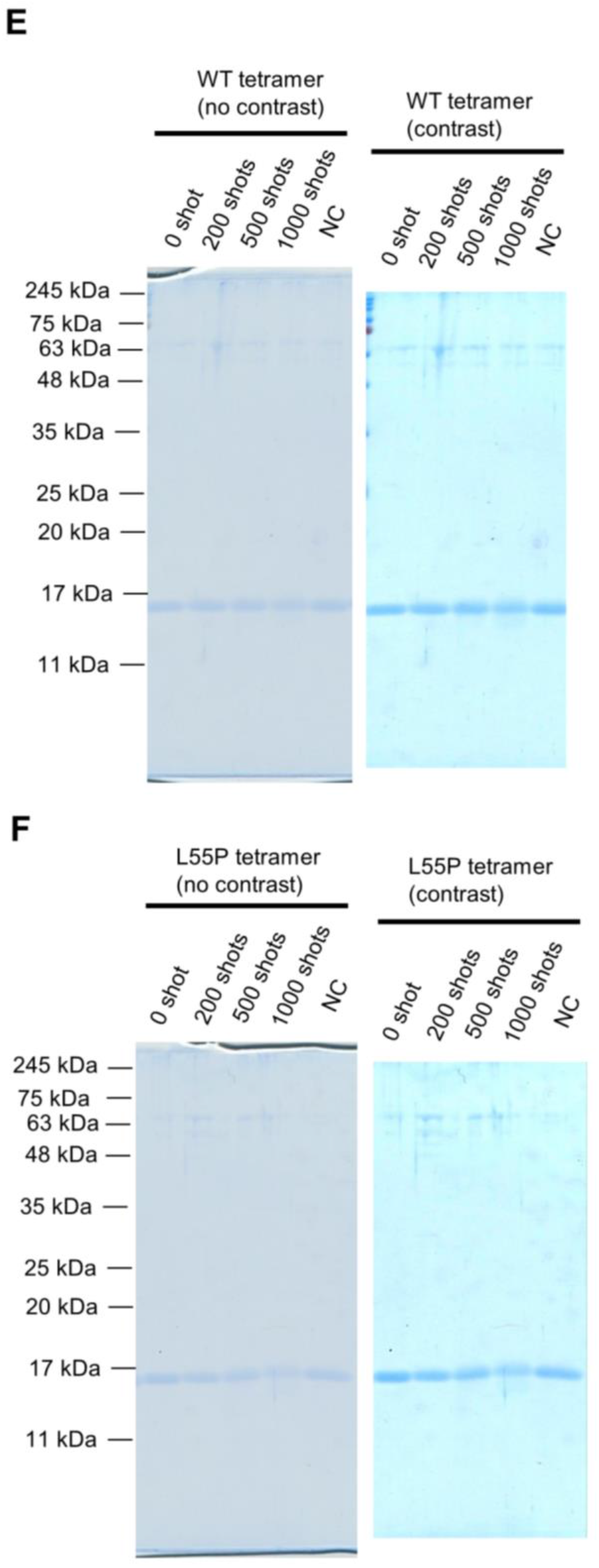
Full photographs of gel electrophoresis in Figure 4. Full SDS-PAGE photographs of WT (**A**) and L55P mutant (**B**) amyloid-derived subunits exposed to 1000 pulses at 1 MV/cm and 2 MV/cm. Full SDS-PAGE photographs of WT (**C**) and L55P mutants. (**D**) Amyloid-derived subunits exposed to several different pulses at 2 MV/cm. Full SDS-PAGE photographs of WT (**E**) and L55P mutants. (**F**) Tetramer-derived subunits exposed to several different pulses at 2 MV/cm. Because some bands were too weak to check without contrast, a contrast version was produced.

## References

1. Robinson, K. R. The responses of cells to electrical fields: A review. J. Cell Biol. 101, 2023–2027 (1985).

2. Chang, F. & Minc, N. Electrochemical Control of Cell and Tissue Polarity. Annu. Rev. Cell Dev. Biol. 30, 317–336 (2014).

3. Levine, Z. A. & Vernier, P. T. Life cycle of an electropore: Field-dependent and field-independent steps in pore creation and annihilation. 236, 27–36 (2010).

4. Perrier, D. L., Rems, L. & Boukany, P. E. Lipid vesicles in pulsed electric fields: Fundamental principles of the membrane response and its biomedical applications. Advances in Colloid and Interface Science 249, 248–271 (2017).

5. Rems, L. & Miklavčič, D. Tutorial: Electroporation of cells in complex materials and tissue. J. Appl. Phys. 119, 201101 (2016).

6. Miklavcic, D., Rols, M.-P., Haberl Meglic, S., Rosazza, C. & Zumbusch, A. Gene Electrotransfer: A Mechanistic Perspective. Curr. Gene Ther. 16, 98–129 (2016).

7. Chen, W. & Lee, R. C. Altered ion channel conductance and ionic selectivity induced by large imposed membrane potential pulse. Biophys. J. 67, 603–612 (1994).

8. Semenov, I., Xiao, S., Kang, D., Schoenbach, K. H. & Pakhomov, A. G. Cell stimulation and calcium mobilization by picosecond electric pulses. Bioelectrochemistry 105, 65–71 (2015).

9. Barzanjeh, S., Salari, V., Tuszynski, J. A., Cifra, M. & Simon, C. Optomechanical proposal for monitoring microtubule mechanical vibrations. Phys. Rev. E 96, (2017).

10. Bezanilla, F. How membrane proteins sense voltage. Nature Reviews Molecular Cell Biology 9, 323–332 (2008).

11. Tuszynski, J. A., Wenger, C., Friesen, D. E. & Preto, J. An overview of sub-cellular mechanisms involved in the action of TTFields. Int. J. Environ. Res. Public Health 13, 1–23 (2016).

12. Carr, L. et al. Calcium-independent disruption of microtubule dynamics by nanosecond pulsed electric fields in U87 human glioblastoma cells. Sci. Rep. 7, 41267 (2017).

13. Dutta, D., Asmar, A. & Stacey, M. Effects of nanosecond pulse electric fields on cellular elasticity. Micron 72, 15–20 (2015).

14. Stacey, M., Fox, P., Buescher, S. & Kolb, J. Nanosecond pulsed electric field induced cytoskeleton, nuclear membrane and telomere damage adversely impact cell survival. Bioelectrochemistry 82, 131–134 (2011).

15. Thompson, G. L., Roth, C., Tolstykh, G., Kuipers, M. & Ibey, B. L. Disruption of the actin cortex contributes to susceptibility of mammalian cells to nanosecond pulsed electric fields. Bioelectromagnetics 35, 262–272 (2014).

16. Beebe, S. J. Considering effects of nanosecond pulsed electric fields on proteins. Bioelectrochemistry 103, 52–59 (2015).

17. Timmons, J. J., Preto, J., Tuszynski, J. A. & Wong, E. T. Tubulin’s response to external electric fields by molecular dynamics simulations. PLoS One 13, e0202141 (2018).

18. English, N. J. & Waldron, C. J. Perspectives on external electric fields in molecular simulation: Progress, prospects and challenges. Phys. Chem. Chem. Phys. 17, 12407–12440 (2015).

19. Marracino, P. et al. Tubulin response to intense nanosecond-scale electric field in molecular dynamics simulation. Sci. Rep. 9, 10477 (2019).

20. Baumketner, A. Electric field as a disaggregating agent for amyloid fibrils. J. Phys. Chem. B 118, 14578–14589 (2014).

21. Pandey, N. K. et al. Disruption of human serum albumin fibrils by a static electric field. J. Phys. D. Appl. Phys. 47, (2014).

22. Pakhomova, O. N. et al. Oxidative effects of nanosecond pulsed electric field exposure in cells and cell-free media. Arch. Biochem. Biophys. 527, 55–64 (2012).

23. Takahashi, M., Handa, A., Yamaguchi, Y., Kodama, R. & Chiba, K. Anodic Oxidative Modification of Egg White for Heat Treatment. J. Agric. Food Chem. 64, 6503–6507 (2016).

24. Zhao, W., Tang, Y., Lu, L., Chen, X. & Li, C. Review: Pulsed Electric Fields Processing of Protein-Based Foods. Food and Bioprocess Technology 7, 114–125 (2014).

25. Wei, Z., Ruijin, Y., Yali, T., Wenbin, Z. & Xiao, H. Investigation of the protein-protein aggregation of egg white proteins under pulsed electric fields. J. Agric. Food Chem. 57, 3571–3577 (2009).

26. Blake, C. & Serpell, L. Synchrotron X-ray studies suggest that the core of the transthyretin amyloid fibril is a continuous β-sheet helix. Structure 4, 989–998 (1996).

27. Powers, E. T., Kelly, J. W., Connelly, S., Fearns, C. & Johnson, S. M. The Transthyretin Amyloidoses: From Delineating the Molecular Mechanism of Aggregation Linked to Pathology to a Regulatory-Agency-Approved Drug. J. Mol. Biol. 421, 185–203 (2012).

28. Kučera, O. & Havelka, D. Mechano-electrical vibrations of microtubules-Link to subcellular morphology. BioSystems 109, 346–355 (2012).

29. Chou, K. C. Low-frequency collective motion in biomacromolecules and its biological functions. Biophys. Chem. 30, 3–48 (1988).

30. Ballestrasse, C. L., Ruggeri, R. T. & Beck, T. R. Calculations of the pH changes produced in body tissue by a spherical stimulation electrode. Ann. Biomed. Eng. 13, 405–424 (1985).

31. Meneses, N., Jaeger, H. & Knorr, D. PH-changes during pulsed electric field treatments - Numerical simulation and in situ impact on polyphenoloxidase inactivation. Innov. Food Sci. Emerg. Technol. 12, 499–504 (2011).

32. Kuhn, A. T. & Chan, C. Y. pH changes at near-electrode surfaces. J. Appl. Electrochem. 13, 189–207 (1983).

33. Germain, P. S., Pell, W. G. & Conway, B. E. Evaluation and origins of the difference between double-layer capacitance behaviour at Au-metal and oxidized Au surfaces. Electrochim. Acta 49, 1775–1788 (2004).

34. Singh, M. B. & Kant, R. Theory for anomalous electric double-layer dynamics in ionic liquids. J. Phys. Chem. C 118, 8766–8774 (2014).

35. Singh, M. B. & Kant, R. Theory of Electric Double Layer Dynamics at Blocking Electrode. (2011).

36. Singh, M. B. & Kant, R. Debye-Falkenhagen dynamics of electric double layer in presence of electrode heterogeneities. J. Electroanal. Chem. 704, 197–207 (2013).

